# Isolation and molecular identification of pectinase producing Aspergillus species from different soil samples of Bhubaneswar regions

**DOI:** 10.1101/837112

**Authors:** Sonali Satapathy, Pabitra Mohan Behera, Dhananjay Kumar Tanty, Shweta Srivastava, Hrudayanath Thatoi, Anshuman Dixit, Santi Lata Sahoo

**Affiliations:** Microbiology Research Laboratory, Post Graduate Department of Botany, Utkal University, Vani Vihar, Bhubaneswar, Odisha, India; Computational Biology and Bioinformatics Lab, Institute of Life Sciences, Nalco Square, Bhubaneswar, Odisha, India; Medicinal and Process Chemistry Division, University Department of Pharmaceutical Sciences, Utkal University, Vani Vihar, Bhubaneswar, Odisha, India; Fragrance & Flavour Development Centre, Ministry of MSME, Govt. of India, Kannauj, Utter Pradesh, India; Department of Biotechnology, North Orissa University, Sri Ram Chandra Vihar, Takatpur, Baripada, Odisha, India

**Author notes:** **Corresponding author** Santi Lata Sahoo. **Co-corresponding author** Anshuman Dixit.

**Keywords:** Aspergillus species, Fungal isolates, Pectinase, Multiple Sequence Alignment, Phylogenetic tree

## Abstract

With the significant improvement of human civilization there is a spur in the urban, rural and industrial development, which has a profound effect on the surrounding natural environment. Increased utilization of natural resources is often associated with accumulation of waste materials whose management is crucial for sustainable development of life. Availability of different microorganisms in the soil facilitates the degradation of wastes through their potential enzymatic activities. Pectinase seems to be one of the important enzymes produced by a wide variety of microorganisms contained in the soil. It is mainly involved in maceration and rotting of plant extracts and debris by hydrolysis of 1,4-alpha glycosidic bonds of de-esterified pectate of plant call wall. In this paper we report molecular identification of some pectinase producing Aspergillus species selected from soil samples of five different zones of Bhubaneswar city using molecular biology and computational techniques. Among fifteen fungal isolates studied from these five zones *Aspergillus parvisclerotigenus* was potent for pectinase production next to *Aspergillus niger* in form of halozone of 0.6 mm. It’s 28S rDNA sequence also had some significant identity (>90%) with different subspecies of Aspergillus. We hope that our findings will helpful in genetic manipulation for improvement of fungal strains of isolates. Again large scale use of the improved Aspergillus strains can degrade plant biomass & diverse industrial wastes which will reduce environmental pollution of capital urban like Bhubaneswar.

## Introduction

Nature provides us a platform with wide variety of micro-organisms which has exceptional potentials for producing useful enzymes with huge commercial viability. The enzyme pectinase has marvelous significance in industrial sector (Dayanand and Patil 2003) with increased relevance to commercial production of pulp & paper (Viikari et al. 2001) secondly food & juice processing (Kashyap et al. 2003). Pectinases are heterogenous group of co-related enzymes which can hydrolyze mostly the pectic substances mainly found in plants. There has been validated proof of pectinolytic enzymes in micro-organisms and higher plants (Jayani et al. 2005). Cell wall extension & plant tissue softening during maturing is often enabled in plants by the help of these enzymes (Ward and Young 1989; Anguilar and Huirton 1990). In addition, this glyco-proteinaceous biomolecule maintains ecological balance through decomposition of plant wastes and recycling them. Plant pathogenicity is one of the major manifestations of pectinolytic enzymes (Singh et al. 1999). As an analysis of the world’s enzyme market, it was noticed that more than 40% of the world’s market share for enzymes were pre-occupied by myco-hydrolytic enzymes and those enzymes are enriching the various industrial processes and technologies (Archer and Peberdy 1997). Microbial pectinases accounts to more than 25% of sales of global food enzymes. Predominantly commercial preparation of pectinase has been reported from fungal species such as *Aspergillus* and *Penicillium* (Silva et al. 2005; Botella et al. 2007). Fungal pectinases which are acidic in nature enable reduction of bitterness & cloudiness of food product especially fruit juices (Patidar et al. 2018) while pectinases which are alkaline in nature have high utility in textile industry for enabling degumming of fiber crops and also the alkaline pectinases have usage in producing high quality papers, extraction of oils, waste water treatment & fermenting tea or coffee (Solbak et al. 2005). Because of their efficacy, pectinases are one of the most important enzymes for usage in industrial sector. Nearly 10% of the total enzyme manufacturing is production of pectinase (Pedrolli et al. 2009). In this competent world, the utilities of fungal enzymes are increasing day by day hence a herculean research work was taken up to perform sequence of events right from isolation, enumeration, identification and screening of innate fungal species to deep dive and analyze the hydrolytic efficiency of fungus for imminent prospective applications across the globe.

## Materials and methods

### Chemicals and media

The chemicals and media used in the present research work were entirely of analytical grade and microbiological grade and were procured from SRL Pvt. Limited, Hi-Media Limited, Merck India Limited and Sigma Chemicals Co. (USA).

### Selection of study site and soil sample collection

Bhubaneswar is the capital city of Odisha, India. It is located in the eastern coastal plains and south-west of the Mahanadi River, which constitutes the northern boundary of it. Experimental fields (5 × 5 m) selected for soil sampling includes soils rich with decomposed fruits and vegetables from 5 different zones of Bhubaneswar such as Unit-IV (Central Bhubaneswar Zone), Chandrasekharpur (Northern Bhubaneswar Zone), Tomando (Southern fringes), Jaydev vihar (Western Bhubaneswar) and Old town area. After site selection, the particular area was scraped out in order to get rid of soil debris and diminutive plants. Aseptically about 1 Kg of soil samples were collected from each study site for soil analysis and microbial analysis. All the collected soil samples were immediately brought to laboratory and stored in refrigerator at 4 °C for further use.

### Edaphic properties of soil samples

Physical parameters of soil such as soil texture, electrical conductivity, water holding capacity and pH were examined. Some chemical parameters of soil like organic carbon, total nitrogen, total phosphorus and total potassium were also evaluated.

### Isolation of microorganisms

The native fungal populations were isolated from soil samples rich with decomposed fruits and vegetables by serial dilution and spread plate method (Mehta et al. 2013). One gram of soil samples from each collection site were pooled and homogenized in sterile distilled water and 10-fold serial dilutions were prepared. 0.1 ml of dilution was spreaded on Potato dextrose agar medium plate amended with cefixime (50 mg/100 ml) and pH values adjusted to 5.6 ± 0.2. The culture plates were incubated under sterile conditions at 30 ± 1 °C for 7 days. The mixed cultures were sub cultured in order to get pure cultures. The fungal isolates thus obtained were stored in refrigerator at 4 °C until further use.

### Screening of isolates for pectinase production

The fungal isolates were screened for biosynthesis of pectinase. This was carried out by inoculating (point inoculation) the fungal isolates on pre-poured PDA plates supplemented with Pectin 10gm/lt and Peptone 5gm/lt as per Hankin and Anagnostakis with certain modifications. The culture plates were incubated under sterile conditions at 30 ± 1°C for 5 days. After complete incubation the plates were flooded with 1% cetyl trimethyl ammonium bromide (CTAB) for 15mins in order to observe the zone of clearance which thus was the indication of extracellular pectinase production (Patil and Chaudhari 2010).

Zone of clearance (Halozone) = Total zone area – Colony diameter

### Identification of pectinase positive fungal isolates

The fungal isolates which indicate the biosynthesis of extracellular pectinase were thus selected and subjected to morphological identification. The selected isolates were identified as per Alexopoulos and Mims (1979), Watanabe (2002) based on their macro and microscopic characteristics following the protocol of Sethi et al. (2013). For micromorphological observations, microscopic mounts were made with lactophenol cotton blue. Phase contrast microscopy was performed with uncoated samples in order to contemplate the shape and size of head, vesicles, conidiophore and conidia, presence of phialides and metulas. For macromorphological observations, fungal isolates were cultured on PDA medium under sterile conditions for 7 days at 30°C in dark to assess fungal growth with low water activity. Macromorphology study of isolates epitomized colony form, pattern, colour, texture, margin, elevation, reverse colour of colony, quantity of aerial hyphae, etc.

### Molecular Identification of the hyperproducer/potent strain

The fungal isolate which performed as the best biosynthesizer of pectinase enzyme was thus subjected to molecular identification. Genomic DNA was isolated from *Aspergillus parvisclerotigenus* SSB9, potent strain by the method of Moller et al. (1992) and confirmed by 1.2% Agarose gel electrophoresis. The resulting DNA was used as a template to amplify the D1/D2 region of LSU (large subunit of 28S rDNA) of ribosomal DNA. The primers used were DF_0616204_ITS1F_G11 and DR_0616204_ITS4_D12. PCR reactions of the 28S rDNA were performed in a final volume of 20 µl containing forward primer-2µl, reverse primer-2 µl, Template DNA-2 µl, PCR master mix-6µl and RNase free water-8µl. DNA primers were used in the forward (DF_0616204_ITS1F_G11) and reverse (DR_0616204_ITS4_D12) directions. The amplifications were carried out in a Thermocycler (Applied Biosystem) and programmed for an initial denaturation of 5 min at 94 °C, followed by 30 cycles of 45 s at 94 °C, 40s at 54 °C and 1 min at 72 °C for extension and annealing and for a final extension of 10 min at 72 °C. The amplified PCR product was purified in a mini column to remove contaminants and sequenced on an Applied Biosystem Instrument Prism 310 Analyzer using primers DF and DR. The sequenced data were compared with the standard sequences in the GenBank (Benson et al. 2011) of NCBI (National Center for Biotechnological Information) using the BLAST (Basic Local Alignment Search Tool) (Altschul et al. 1990). Species were identified based on the percentage similarity with the known species sequences in the data base. Electrophoresis of the PCR amlicon was performed in 1.2 5% (w/v) agarose gels in TBE buffer (89 mM Tris-HCl, 89 mM boric acid, 2 mM EDTA; pH 8.0), then stained with ethidium bromide and the developed gel was visualized under UV transilluminator. A consensus sequence of 601 bp of D1/D2 region of 28S rDNA gene was generated from forward and reverse sequence data using aligner software. The resulting consensus sequence was submitted to NCBI with the accession number KX928754.

### *In Silico* Molecular phylogenetics analysis

Prior to phylogenetic analysis the sequence (KX928754) was opened in BioEdit (Hall et al. 1999) for analysis of genetic features. It has a molecular weight of 182987.00 Daltons with more G+C content (57.74%) than A+T (42.26%) content. Then the sequence was queried in NCBI BLASTn for fetching of highly similar sequences with megablast algorithm. About 50 similar sequences were selected from the result for construction of a distance based phylogenetic tree using Neighbor Joining (NJ) (Saitou et al. 1987) method with maximum sequence difference of 0.75. A circular layout of the tree was generated and sorted by number of children.

## RESULTS

### Edaphic properties of soil sample

The strata of surface soil can be contemplated as an eventful pocket that stores a wide array of soil dwelling microorganisms and catalyzes all essential microbial processes. The physical and chemical properties of 5 different soil samples obtained from different zones of Bhubaneswar were analyzed. The soil from Chandrasekharpur area was having alkaline pH value i.e. 8.30 followed by Old town (7.47) and slightly acidic in Unit-IV area i.e. 5.98. The organic carbon content was highest in soil of Tomando area i.e. 0.95% and least in Chandrasekharpur area (0.34%). The water holding capacity was almost same in soil of Chandrasekharpur and Jaydev vihar area but highest in Old town area (Table 1). The total nitrogen and phosphorus content was found to be 387 Kg/ha and 76 Kg/ha in Chandrasekharpur area soil which was highest amongst all. Similarly total potassium content in Chandrasekharpur and Tomando area were 238 Kg/ha and 202 Kg/ha followed by Old town and Unit-IV area i.e. 487 Kg/ha, 450 Kg/ha and least was obtained in Jaydev vihar area which was 149 Kg/ha.

**Table 1:** Physical and chemical properties of different soils collected for isolation of microorganisms

| Sl. No. | Soil Samples | pH | E.C. (dS/m) | Organic C (%) | Available N (Kg/ha) | Available P (Kg/ha) | Available K (Kg/ha) | WHC (%) |
| --- | --- | --- | --- | --- | --- | --- | --- | --- |
| 1 | Chandrasekharpur | 8.30 | 0.24 | 0.34 | 387 | 76 | 238 | 29.65 |
| 2 | Tomando | 6.85 | 0.16 | 0.95 | 224 | 67 | 202 | 28.25 |
| 3 | Old Town | 7.47 | 0.23 | 0.49 | 341 | 59 | 487 | 31.92 |
| 4 | Unit- IV | 5.98 | 0.10 | 0.74 | 312 | 61 | 450 | 30.52 |
| 5 | Jaydev Vihar | 7.10 | 0.10 | 0.65 | 236 | 25 | 149 | 29.62 |

### Isolation of fungal isolates

For obtaining the desired microbe from the soil samples of different areas adjacent to Bhubaneswar, serial dilution and spread plating isolation techniques were used. Subsequently, the fungal isolates were sub cultured into their respective growth media until pure cultures were isolated (Fig 1). A total of 354 fungal cultures were obtained from soil samples rich with decomposed fruits & vegetables of 5 different areas of Bhubaneswar. Highest numbers of fungi were isolated from Unit-IV area (13 × 10^8^ CFU/g) followed by Tomando area (5 × 10^8^ CFU/g), Chandrasekharpur area (4 × 10^8^ CFU/g) and least fungal isolates were obtained from Old Town area (2 × 10^8^ CFU/g), Jaydev vihar area (3 × 10^8^ CFU/g).

**Fig 1:**
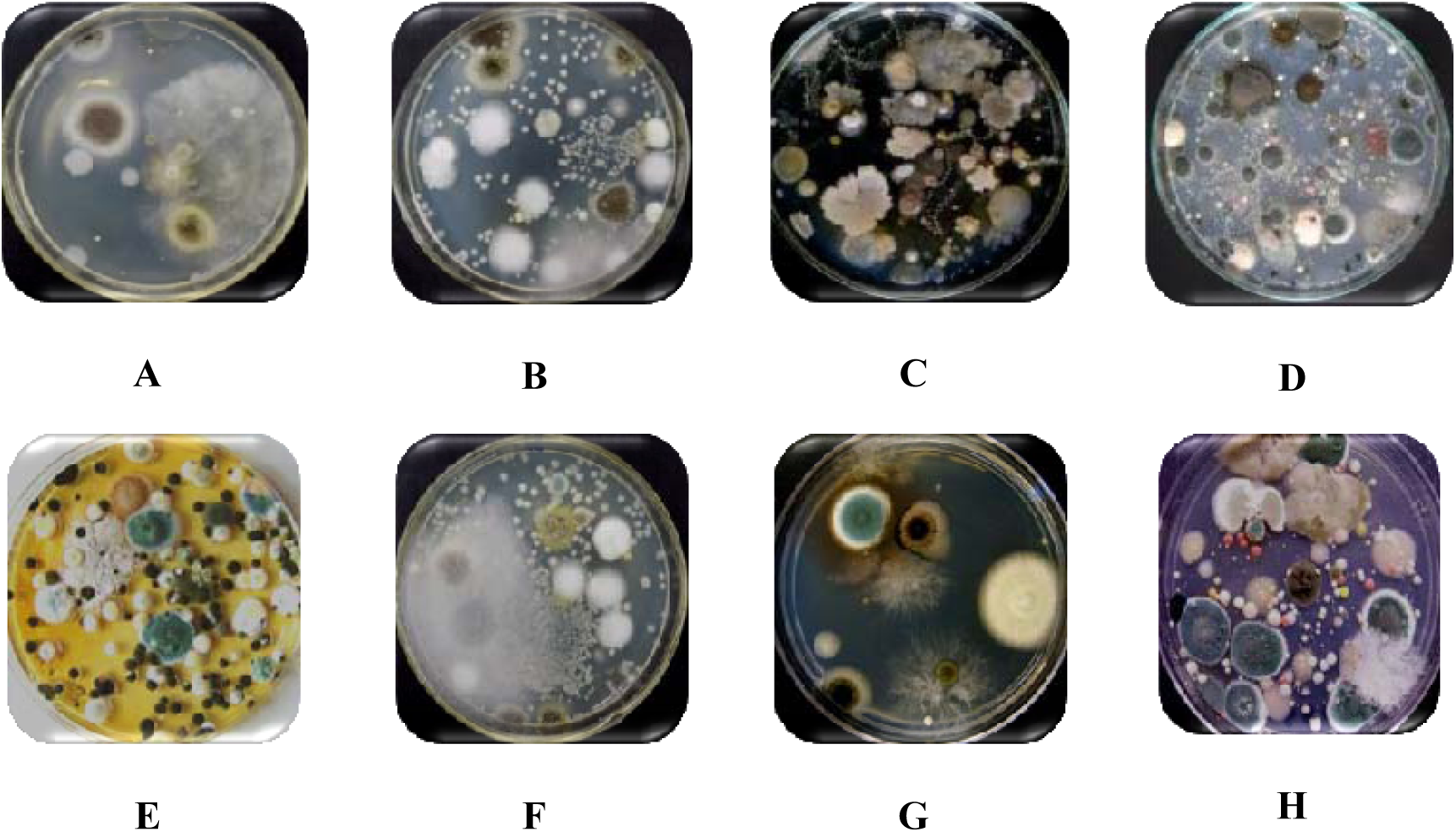
Mixed cultures isolated from different soil samples

### Screening of isolates for pectinase production

The fungal cultures obtained from soil samples of 5 different localities of Bhubaneswar were screened for the ability to biosynthesize extracellular pectinase. About 15 fungal isolates were competent enough to biosynthesize extracellular pectinase (Fig 2). Among those isolates, the genus *Aspergillus* was the predominant one. The maximum zone of clearance was observed in fungal isolate PDI 8 *Aspergillus niger* (Table 2, Fig 3) followed by PDI 13 *Aspergillus parvisclerotigenus*. As a lot of exploration on pectinase production by *Aspergillus niger* has already been accomplished so the subordinate strain PDI 13 *Aspergillus parvisclerotigenus* was thus selected for further studies. This strain can be a prospective candidate for the economic production of high-valued products using different fermentation techniques.

**Table 2:** Qualitative assay of pectinase from fungal isolates (Plate Assay)

| Sl. No. | Fungal Isolates | Halozone (cm) |
| --- | --- | --- |
| 1 | PDI 1 | 0.5 |
| 2 | PDI 2 | 0.4 |
| 3 | PDI 3 | 0.5 |
| 4 | PDI 4 | 0.2 |
| 5 | PDI 5 | 0.1 |
| 6 | PDI 6 | 0.3 |
| 7 | PDI 7 | 0.4 |
| 8 | PDI 8 | 0.8 |
| 9 | PDI 9 | 0.1 |
| 10 | PDI 10 | 0.2 |
| 11 | PDI 11 | 0.1 |
| 12 | PDI 12 | 0.4 |
| 13 | PDI 13 | 0.6 |
| 14 | PDI 14 | 0.3 |
| 15 | PDI 15 | 0.3 |

**Fig 2:**
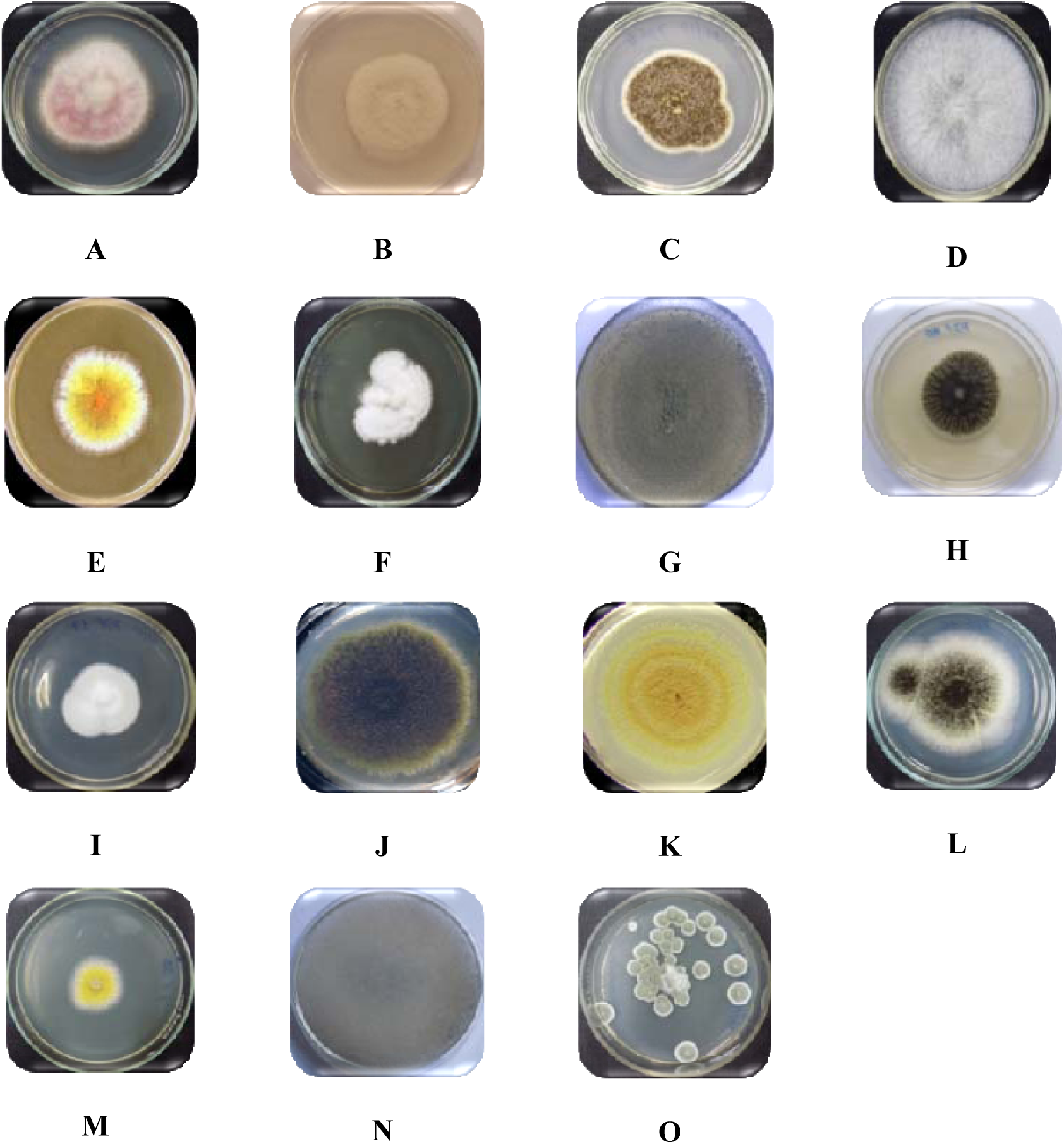
Pure culture plates of pectinase positive fungi. (A: *Fusarium oxysporum*, B: *Aspergillus candidus*, C: *Asperillus* sp., D: *Mucor* sp., E: *Aspergillus glaucus*, F: *Aspergillus thermomutans*, G: *Aspergillus flavus*, H: *Aspergillus niger*, I: *Fusarium semitectum*, J: *Aspergillus nidulans*, K: *Asperrgillus tamari*, L: *Aspergillus awamori*, M: *Aspergillus parvisclerotigenus*, N: *Mucor hiemalis*, O: *Aspergillus fumigatus*)

**Fig 3:**
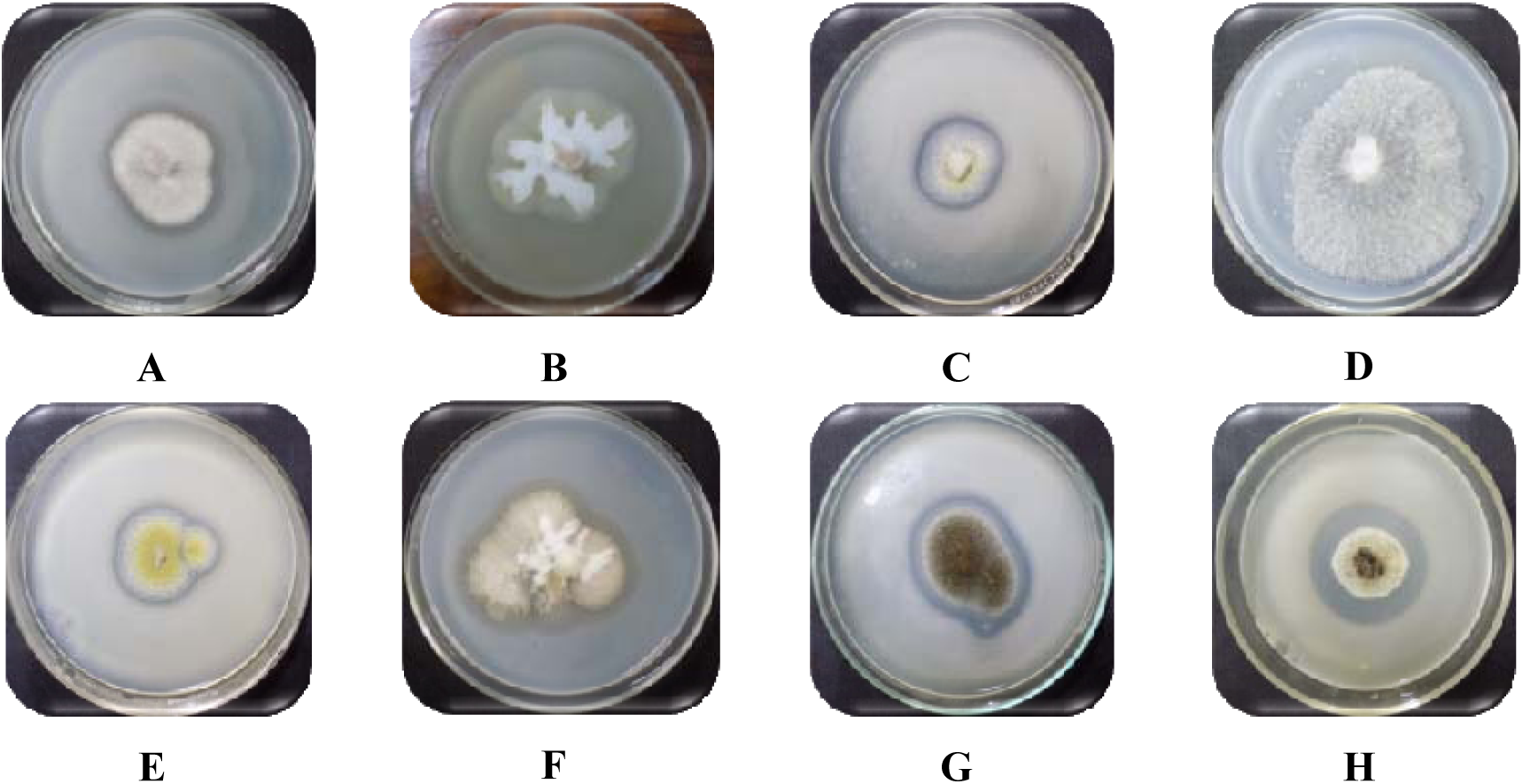
Screening of fungal isolates for pectinase production. (A: PDI 1, B: PDI 9, C: PDI 7, D: PDI 4, E: PDI 13, F: PDI 2, G: PDI 3, H: PDI 12)

### Identification of pectinase positive fungal isolates

The selected fungal isolates were identified as per Alexopoulos and Mims (1979) and Watanabe (2002) based on their macro and microscopic characteristics and the protocol developed by Sethi *et al.* (2013). In all the five different soil samples, the genus *Aspergillus* was the predominant one. A total of 15 fungal isolates exhibited positive receptivity for extracellular pectinase biosynthesis (Fig 3). Out of which 11 fungal isolates represented the efficacious genus *Aspergillus* (Table 3, Fig 4) with species varying from *Aspergillus candidus, Aspergillus flavus, Aspergillus glaucus, Aspergillus niger, Aspergillus parvisclerotigenus, etc.* There were two isolates from genus *Fusarium* (*Fusarium oxysporum* and *Fusarium semitectum*) and two isolates from the genus *Mucor* (*Mucor hiemalis* and *Mucor sp*.). The observation made validated their affiliation up to genus level, not sufficient enough for the identification up to the species level. Hence a specific identification is also required, which is underway.

**Table 3:** Macro and micro-morphological features of pectinase positive fungal isolates

| Sl. No. | Name of the fungus | Macroscopic characteristics | Microscopic characteristics |
| --- | --- | --- | --- |
| 1 | *Fusarium oxysporum* | Surface colony colour is pinkish white. Reverse plate colour is white to slight red. Raised colony with entire margin and cottony texture. | Conidiophores are hyaline, simple, short or not well differentiated from hyphae, bearing masses of spore at the apexes. Conidia are of two kinds: macroconidia and microconidia Macroconidia are boat-shaped, with slightly tapering apical cells and hooked basal cells. The size of the macroconidia ranged from 20 - 30 $\mu\text{m}$ in length and 5.00 - 6.75 $\mu\text{m}$ in breadth. Microconidia are formed singly, oval to reniform and without any septation. The size of microconidia ranged from 7 -8 $\mu\text{m}$ in length and 2 - 3 $\mu\text{m}$ in breadth. Chlamydospores are brown, globose, and usually solitary. |
| 2 | *Aspergillus candidus* | Surface colony colour is white. Reverse plate colour is off white. Colony is flat and texture is a velvety to powdery. | Conidiophores are hyaline, simple (300 - 350 $\mu\text{m}$ long), thin cell wall, inflated ellipsoidally at the apex forming often noded vesicle of pale brown colour. Phialides are uniseriate. Conidia are phialosporous, pale green, globose (3.0 - 4.5 $\mu\text{m}$ in diameter), slightly echinulate. |
| 3 | *Aspergillus sp.* | Surface colony colour is brownish black. Reverse plate colour is white to creamy. Colony is having filiform margin with texture powdery and a bit cottony. | Conidiophores are erect (1.0 - 1.5 mm long), simple or rarely branched, smooth in the surface, bearing dark bean-shaped conidial heads composed of densely catenulate conidia borne on uniseriate phialides developed on elliptical vesicles. Conidia are phialosporous, globose to subglobose (3.0 - 3.5 $\mu\text{m}$ in diameter) and rough in the surface. |
| 4 | *Mucor sp.* | Surface colony colour is white. Reverse plate colour is creamy white. Uniformly raised colony with entire margin and texture cottony or wooly. | Wide hyphae approx 6 - 15 $\mu$. Sporangiphores are long, hyaline, erect, branched sympodically, bearing sporangia terminally, tapering gradually from base towards apex, rhizoidal, ellipsoidal. Sporangia are about 250 $\mu\text{m}$ in diameter, terminal, black, columellate, bearing sporangiospores of about 12 $\mu\text{m}$ in length and 100 - 180 $\mu\text{m}$ in diameter. Collumellae are globose or sub globose. |
| 5 | *Aspergillus glaucus* | Surface colony colour is orange green with yellow areas. Reverse plate colour is yellowish to brown. Colony is circular, flat and have entire margin. | Conidiophores are erect (200 - 350 $\mu$ long, 7 - 12 $\mu$ wide) having round to columnar vesicle of 15 – 30 $\mu$ diameter. Conidia are phialosporous, subglobose to slightly echinulate (5.0- 5.5 $\mu\text{m}$ in diameter), borne on uniseriate phialides. |
| 6 | *Aspergillus thermomutans* | Surface colony colour is white. Reverse plate colour is off white. Colony is slightly raised with entire margin and surface is smooth and velvety. | Vesicles are small (10 - 12 $\mu$), round, loosely, radiating head. Conidia are smooth, short (2.0 - 2.5 $\mu\text{m}$ in diameter), colourless borne on biseriate phialides and forms long chains. Primary phialides shorter than secondary phialides. |
| 7 | *Aspergillus flavus* | Surface colony colour is yellow to green. Reverse plate colour is goldish to red brown. Colony texture is powdery. | Conidiophores are heavy walled, uncoloured, stipes are hyaline and coarsely roughened, usually less than 1 mm in length. Vesicles are subglobose or globose, varying from 10 to 55 μm in diameter. Phialides are uniseriate or biseriate. The primary branches are upto 8 mm in length, and these secondary upto 4 mm in length. Conidia are globose to subglobose (3 to 6 μm in diameter), conspicuously echinulate, pale green in colour. |
| 8 | *Aspergillus niger* | Surface colour is homogeneous black. Reverse plate colour is white. Uniformly raised colony with entire margin, regular to elliptical colony and surface is wrinkled and powdery. | Conidiophore are hyaline or pale brown, erect (1.5 - 2 mm long), simple, thick walled, with foot cells basally, inflated at the apex. Vesicles are globose (50 μm in diameter) bearing conidial heads composed of catenulate conidia. Phialides (20 - 30 μm) are uniseriate or biseriate, acutely tapered at apex. Conidia are phialosporous, brown, black in mass, globose (4 - 5 μm in diameter), minutely echinulate. |
| 9 | *Fusarium semitectum* | Surface colour is pure white. Reverse plate colour is slight yellowish white. Umbonate colony with entire margin and velvety texture. | Conidiophores are hyaline, simple, short or indistinct from normal hyphae, bearing spore masses at the apex. Conidia are phialosporous, hyaline and of two kinds: macroconidia and microconidia. Macroconidia are abundant and borne in the aerial mycelium, mostly straight or slightly curved, 3 - 5 septate, measuring about 35.0 to 45.5 $\mu\text{m}$ in length. Microconidia were 1 - 2 septate. |
| 10 | *Aspergillus nidulans* | Surface colony colour is green, buff to yellow. Reverse plate colour is dull yellow. Uniformly raised colony with entire margin, circular form and hairy to powdery texture. | Conidiophores are 70 - 150 $\mu$ long and 3 - 6 $\mu$ wide. Vesicles are hemispherical of about 8 - 12 $\mu$ diameter. Phialides are found on distal portion of vesicle and are biserial. Conidia are round or spherical of 3 - 4 $\mu$ in diameter, smooth and brown in colour. |
| 11 | *Aspergillus tamari* | Surface colony colour is mustard yellow. Reverse plate colour is pale yellow. Colony is having curled margin with rough and powdery texture. | Conidiophores are 1 - 2 mm in length, hyaline, roughened. Vesicles are spherical of about 10 - 50 $\mu\text{m}$. Phialides are uniseriate and biserial, covering the entire surface of the vesicle. Conidia are echinulate to tuberculate, subspherical (5 - 8 $\mu\text{m}$ in diameter). |
| 12 | *Aspergillus awamori* | Surface colony colour is blackish brown with off white border. Reverse plate colour is faint yellow. Raised colony with entire margin, circular form and cottony to powdery texture. | Conidiophores are aseptate, straight, mono to bi-layer, usually arose from the foot cell of basal mycelium; stipes straight, hyaline to pale brown, not constricted below the vesicles. Vesicles are hemispherical to elongate (43 - 54 $\mu\text{m}$ in diameter). Conidia are globose to subglobose (3.6 - 4.8 $\mu\text{m}$ in diameter), rough, grey brown, parallel in chains, borne on compact uniseriate phialides. |
| 13 | *Aspergillus parvisclerotigenus* | Surface colony colour is greenish yellow with whitish border. Reverse plate colour is creamy white. Circular form colony with entire margin and powdery to dusty texture. | Conidiophores are thick walled, uncoloured, roughened, usually less than 1.5 mm in length. Vesicles are becoming subglobose or globose, varying from 10 to 45 mm in diameter. Phialides are uniseriate or biseriate. Conidia are phialosporous, greenish yellow, globose to subglobose (3 - 5 mm diameter). |
| 14 | *Mucor hiemalis* | Surface colony colour is ash white. Reverse plate colour is slight yellowish. Raised colony with texture cottony or wooly. | Sporangiophores are erect, pale, reddish yellow, simple or branched, bearing sporangia terminally. Sporangia are globose (230 - 250 µm in diameter), yellow, smooth, columellate on dehiscence. Collumellae are globose. Sporangiospores are long (12 µm in length, 150 - 275 µm in diameter). |
| 15 | *Aspergillus fumigatus* | Surface colony colour is dark green to grey. Reverse plate colour is white to tan. Raised colony with entire margin, circular form and smooth velvety texture. | Conidial heads are columnar and uniseriate. Conidiophores are short, smooth-walled and have conical-shaped terminal vesicles which support a single row of phialides. Conidia forms long chains, globose to subglobose (2.5 - 3.5 µm in diameter), green and finely roughened. |

**Fig 4:**
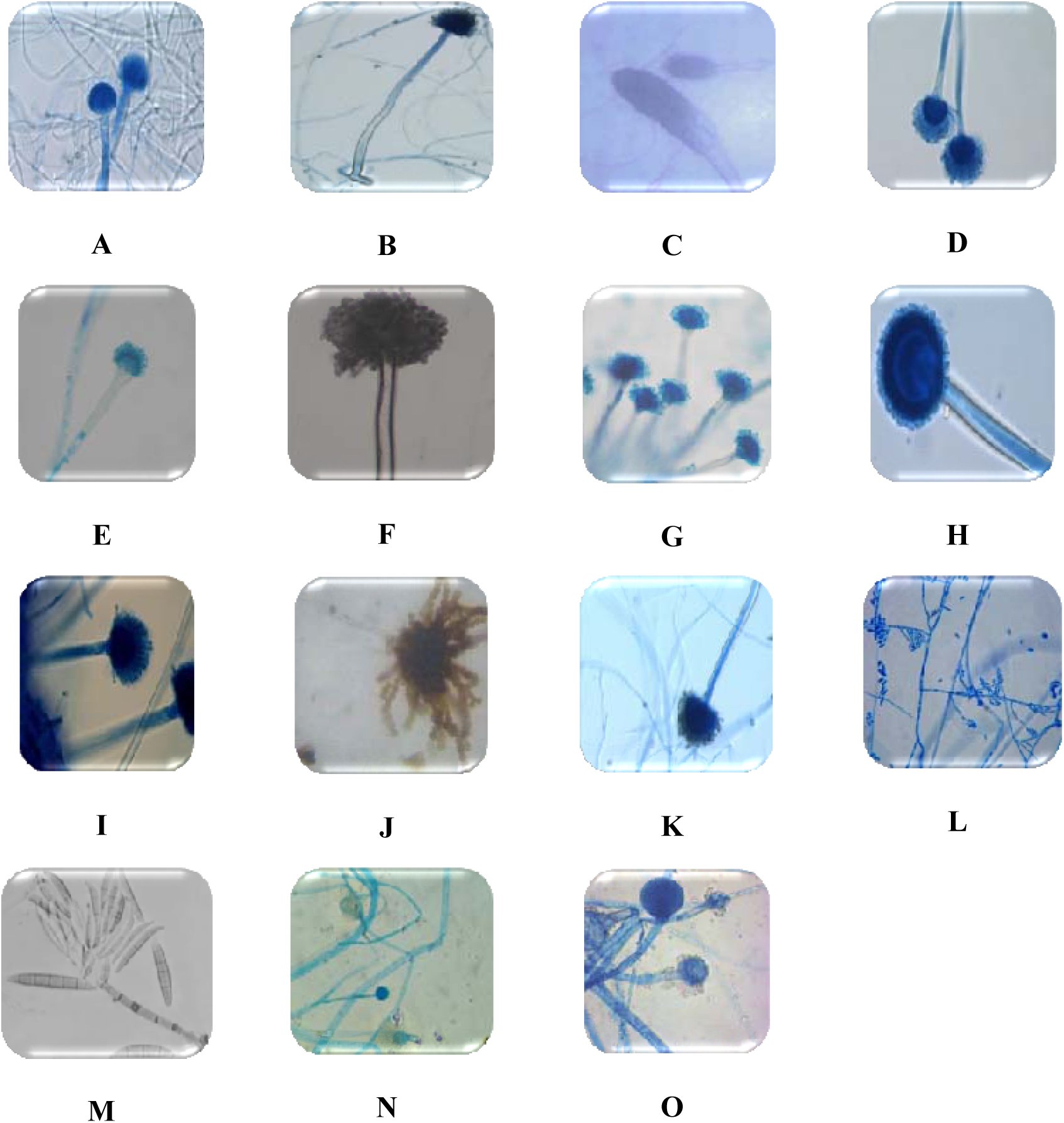
Phase contrast microscopic view of pectinase postive fungi. (A: *Aspergillus awamori*, B: *Aspergillus candidus*, C: *Aspergillus clavatus*, D: *Aspergillus flavus*, E: *Aspergillus fumigatus*, F: *Aspergillus glaucus*, G: *Aspergillus nidulans*, H: *Aspergillus niger*, I: *Aspergillus parvisclerotigenus*, J: *Aspergillus tamari*, K: *Aspergillus thermomutans* L: *Fusarium oxysporum*, M: *Fusarium semitectum*, N: *Mucor hiemalis*, O: *Mucor* sp.)

### *In Silico* Molecular phylogenetics analysis

From the phylogenetic analysis of the designed NJ tree shown in Fig. 5 it was evident that the input Query_16931 is identical (100%) with sequence KX928754 which is the same sequence being deposited by the authors in NCBI GenBank on 30-SEP-2016. Again out of the fifty sequences including the Query_16931 none of the sequences were left for the construction of NJ tree. This may be due to selection of Max. Seq. Difference value = 0.75 which indicates the maximum allowed fraction of mismatched bases in the aligned region of any pair of sequences is less than 0.75. The immediate neighbors of KX928754 were KP278179 and HQ285580 categorized as *Aspergillus oryzae*. Among fifty sequences we were happy to find KC964101, a 1135 bp long linear DNA categorized as *Aspergillus parvisclerotigenus*. The multiple sequence alignment of the query and four described sequences is shown in Fig. 6.

**Fig 5:**
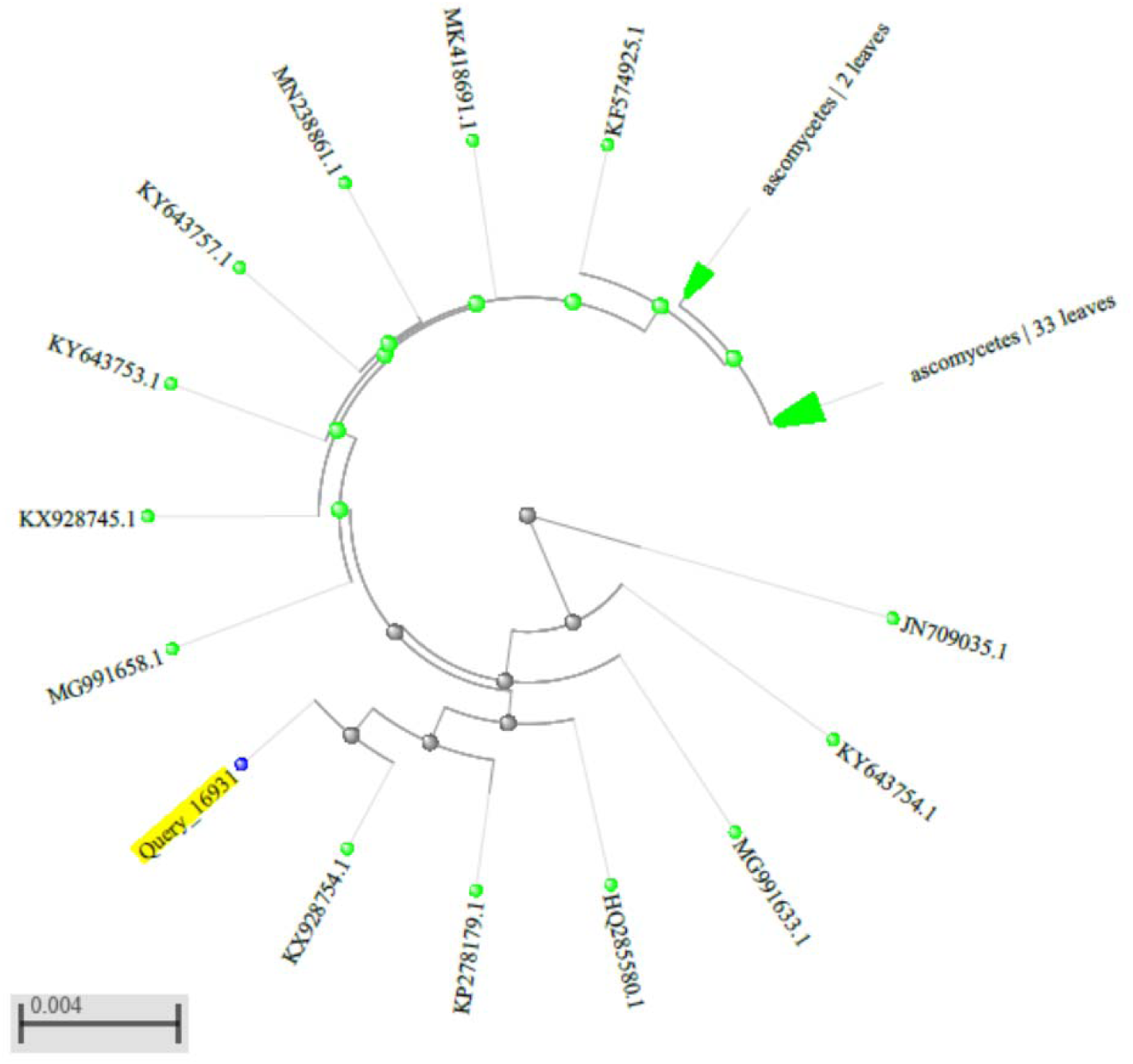
The distance based NJ phylogenetic tree featuring the Query_16931 (highlighted in yellow) with all other highly similar sequences.

**Fig 6:**
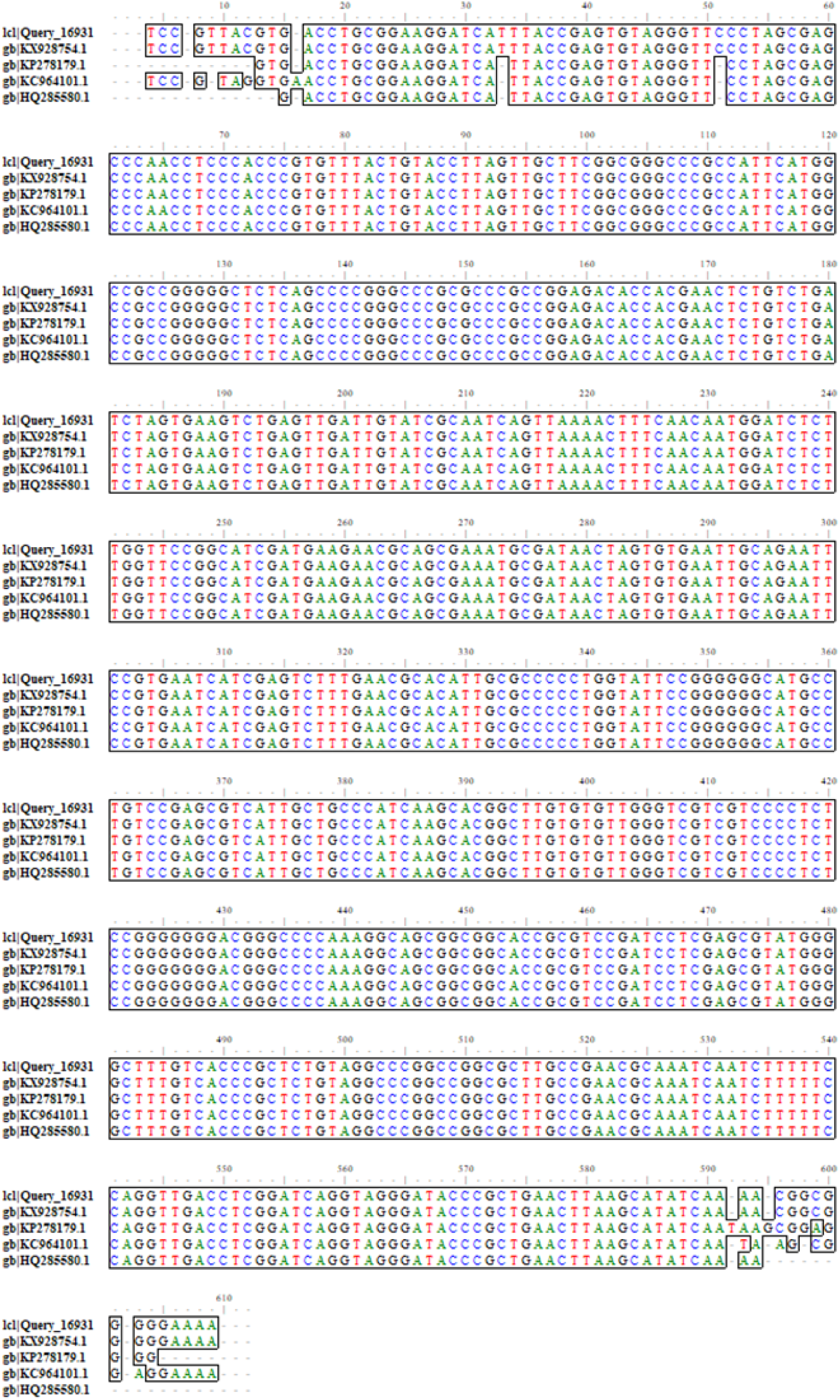
The multiple sequence alignment of Query_16931 and other four sequences showing the identical bases among the five sequences highlighted in outline and coloured by the type of bases.

## Discussions

In this contemporary world researchers have seen the micro-organisms to be one of the leading sources of novel industrial enzymes which invigorate attention towards reconnaissance of extracellular enzymatic biosynthesis in various microorganisms (Jayani et al. 2005). In this experiment, potential pectinase producing microorganisms were isolated from soil using serial dilution and streak plating techniques. Similar analogous processes/techniques were conducted on fungi isolation from soil samples (Reddy and Sreeramulu 2012). Fungi form a co-productive symbiotic network in presence of all the natural media like soil, water and air. Another majestic creation of nature is the marvelous thing called “soil” which is the repository bank for various living organisms that world has seen, felt or interacted. Soil provides a platform for creating, nourishing and improvising living organisms. For fungi, soil is one of the primary habitats that enable its growth and development. Researches fortify the very phenomenal fact of micro-organisms symbiotic relationship with the living world and their significant contribution in enriching the habitat of human beings by conventional industrial microbiology, advanced biotechnology (Recombinant DNA Technology) and next generation industrial fermentation processes. Pectinase producing microorganisms can be manipulated genetically & environmentally for strain improvement and augmenting yield (Bhardwaj and Garg 2014). Myco-hydrolytic enzymes contributes about 40% of the world’s market share for enzymes and those enzymes are ecofriendly and imperishable in nature with wide range of industrial applications. High utility applications of pectinase enzymes provide strong urge to screen micro-organisms that produces pectinase & has more novel properties with increased enzyme activities and with significant production.

In this research work, the total fungal isolates from different soil samples were screened for extracellular biosynthesis of pectinase. Only 15 fungal isolates were ear-marked as the pectinase producers, hence subjected for morphological identification tentatively. The genus *Aspergillus* was the predominant one among all (Table 4) with *Aspergillus niger* and *Aspergillus parvisclerotigenus*. As a lot of exploration on pectinase production by *Aspergillus niger* has already been accomplished but exploiting the propitious isolate *Aspergillus parvisclerotigenus* for pectinase biosynthesis has not been reported so far. The computational analysis citing selection of true homologues of this potential strain through database similarity search, their multiple sequence alignment and subsequent phylogeny prediction also suggested that it was identical (98.50%) to a reported strain of *Aspergillus parvisclerotigenus* KC964101 (Adjovi et al. 2014) in GenBank. The alignment of these potential strains and two other homologues strains shown in Fig.7 also suggests the conservation of nucleotides among the strains with same genetic and subsequent physical expression characteristics.

Hence this strain can be a prospective candidate for the economic production of high-valued products by amalgamation of different fermentation techniques and genetic engineering approaches which not only can lower down the processing cost related to enzyme biosynthesis but also could ramp up its novel industrial applications. Stupendous analogous work (Reddy and Sreeramulu 2012; Anisha et al. 2013) has been done around the globe for validating the high utility usage of the genus *Aspergillus* in generation of pectinolytic enzymes.

## Conclusion

The immacular contribution of Aspergilli to this stupendous society paves the way for an inevitable concrete conclusion that they are exquisitely marvelous in comparison to other enzyme producers. In addition to that they have excellent value-added capabilities to biosynthesize heterologous protein (Dalboge 1997). Therefore, this breadth & depth of research work will enunciate a path breaking innovation in finding ways of genetic manipulation for strain improvement of isolates. Aspergilli with manipulation have brilliant properties for enhancing sustainability of human society as they can degrade plant biomass & diverse industrial wastes which will reduce environmental pollution.

